# The adaptive function of sexual reproduction: resampling the genotype pool

**DOI:** 10.1101/2020.06.14.151001

**Authors:** Donal A. Hickey, G. Brian Golding

## Abstract

Recombination allows a finite population to resample the genotype pool, i.e., the universe of all possible genotypic combinations. This is important in populations that contain abundant genetic variation because, in such populations, the number of potential genotypes is much larger than the number of individuals in the population. Here, we show how recombination, in combination with natural selection, enables an evolving sexual population to replace existing genotypes with new, higher-fitness genotypic combinations. In contrast to this, an asexual population is limited to selection among existing genotypes. Since it has been shown that most eukaryotic species are genetically polymorphic, our model can explain the ubiquity of sex among such species. The model also indicates that classic population genetics theory is applicable to ecological studies of natural selection acting on standing genetic variation.

## Introduction

The function of sexual reproduction has been the subject of much debate among biologists (see Otto 2009, Meirmans and Strand 2010, Colegrave 2012, for recent reviews). Here, we illustrate the effects of sexual reproduction in a finite population where many genes are undergoing selection. An example of this situation would be where adaptation is based on the standing genetic variation in a local population (Barrett and Schluter 2007, Burke et al 2010, Kosheleva and Desai 2017, McDonald et al 2016, Teotonio et al 2009). One of the earliest explanations for the function of sex was that it combines different beneficial mutations in a single individual, resulting in fitter genotypic combinations (Fisher 1930, Muller 1932). Those authors focused on new beneficial mutations rather than on the standing genetic variation. Maynard Smith (1978) pointed out that, given a low mutation rate, small populations might not contain multiple mutations at the same time. By focusing on the abundant standing genetic variation rather than only new mutations, we avoid Maynard Smith’s concern. Of course, these new mutations will eventually contribute to the standing genetic variation once they reach significant frequencies.

Biologists are well aware that Mendelian genetics in an outbreeding sexual population is essentially a game of chance. In this study, we use a lottery analogy in order to explain, in simple terms, the biological function of genetic recombination in a sexually reproducing population. We point out that in the case of inheritance, the lottery is biased by selectively-induced, heritable changes in the frequencies of individual genetic variants. We argue that it is the interaction between random recombination and deterministic natural selection that allows sexual populations to evolve much more efficiently than asexual populations. This provides a consistent, short-term selective advantage for sexual reproduction.

The genetic lottery is special in a number of ways, and by taking these special features into account we can better understand its function. The first feature to be taken into account is that the winning combination, or optimal genotype, does not change drastically and unpredictably from one generation to the next. Environments do change, but this change usually happens gradually over many generations. Thus, the current environment is a reasonable estimator of environmental conditions in the following generations. The second point to consider is that the number of possible genotypic combinations can be many orders of magnitude larger than the total population size (see discussion below). This means that the chance that even a single individual within the entire population will have the optimal genotype is extremely remote. The third point is that both genetic lotteries and many conventional lotteries can have partial winners even in the absence of an overall jackpot winner. In both cases there is an advantage in having at least some of the winning numbers. Finally, the crucial difference between a conventional lottery and the genetic lottery is that inheritance provides an informational feedback each generation regarding the “winning” genotypic combination. We explain this point in more detail below.

## The model

Students of genetics are familiar with the process of generating various genotypic combinations of alleles at two, three or four variable loci. But we rarely stop to think how this plays out at the genomic level. Eukaryotic genomes are estimated to contain approximately 20,000 genetic loci and many of these loci are genetically polymorphic (Lewontin and Hubby 1966). This discovery of abundant genetic variability in natural populations means that the impact of recombination is much greater than what was previously envisaged (Stearns 1985). If we take the overly conservative estimate that only 1% of the gene loci within the genome are genetically variable, we are still left with 200 genes. If we make a further conservative assumption that none of the genes has more than two allelic variants, we can calculate that there are 2^200^ possible genotypic combinations. This number is larger than 10^60^ which in turn is several orders of magnitude larger than the estimated number of atoms in the solar system. Obviously, no biological population is large enough to contain even a single individual representing each of these combinations. In fact, real populations, because of their finite size, contain only a tiny fraction of all possible genotypic combinations.

The relationship between the number of genetically variable genes and the total number of possible genotypic combinations was first noted by East (1918). He pointed out that ten variable genes could generate 1,024 different genotypes and that twenty variable genes could generate 1,032,576 genotypic combinations. Here, we extend this calculation up to 200 genes. The relationship between the number of variable genes and the corresponding number of genotypes is shown in Figure 1. From the figure it can be seen that once we reach more than thirty genetic loci, the number of possible genetic combinations already exceeds most real population sizes, especially those of macroscopic plants and animals. For example, 30 biallelic loci can generate more than one billion different genotypes. The number of genotypes corresponding to 100 loci is approximately 10^30^, which is larger than the number of atoms in the solar system; the number corresponding to 200 loci is many orders of magnitude larger than this again.

**Figure 1.**
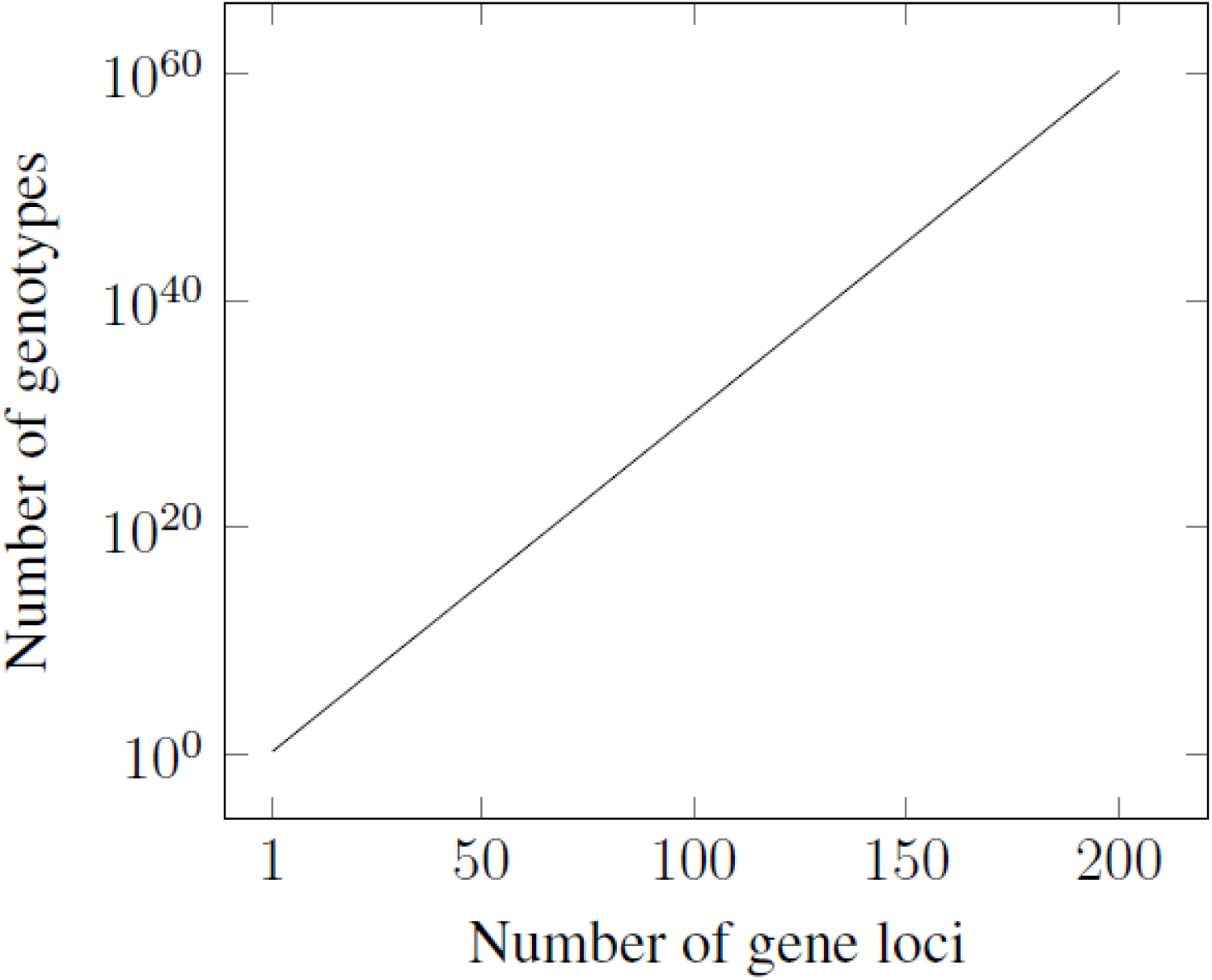
The relationship between the number of genetically variable genes and the number of possible genotypic combinations.

The graph shows the results for up to 200 biallelic loci. The Y axis is on a log scale.

Using the lottery analogy, we can think of an individual genotype containing 200 variable genes as a “ticket” with a list of 200 numbers. In the simple case where we consider only two variants per gene, we can represent the variants as a series of “zeros” and “ones”, where zero represents the more common allelic variant and the number one represents a rarer advantageous variant. Again, for simplification let’s assume that the initial frequency of the advantageous variant at each genetic locus is 0.1 or ten percent. If the tickets are chosen randomly there is a ten percent chance of observing a “1” at any one of the 200 positions. A lottery jackpot winner would have a “1” at all two hundred positions. But the chance of finding such a ticket in the initial population is only 1 out of 10^200^. In other words, we would need to have a number of tickets that is many orders of magnitude larger than the number of atoms in the observable universe in order to have a reasonable chance of observing one such ticket. This means that, in practice, such a combination would not be observed. In fact, all of the tickets would have relatively few ones and many zeros. For any given ticket, we expected to see an average of 20 ones and 180 zeros; this expectation is based on the initial frequencies of ones and zeros. But if the numbers are randomly assigned based on these frequencies, not every ticket will have exactly 20 ones. The numbers will be binomially distributed around the average of 20 and the majority of the values will fall between 12 and 28. This means that there is not only an essentially zero chance of observing a ticket with 200 ones (the jackpot winner), there is also virtually zero chance of observing a ticket with even half that number of ones. So, at first glace, this genetic lottery seems entirely hopeless.

But, as stated above, the genetic lottery is special in that there is a differential survival of tickets depending on their genetic fitness, i.e., the number of “ones” on the ticket. This means that, after selection, tickets containing a higher number of ones will be more represented than those tickets containing less ones. This is the process of natural selection. But natural selection alone can only result in a population containing tickets with the highest number of ones that already exist in the initial population. In our example, this would be about 30 out of the maximum of 200. But if the rounds of selection alternate with rounds of sexual outbreeding and recombination, we observe a further effect. Specifically, as the frequency of “ones” increases due to selection, recombination will randomly recombine them to former a higher average number of “ones” per ticket. Thus, the power of random recombination lies in its ability to adjust the number of “ones” per ticket to match the current frequencies at individual positions. For example, when the frequency of “ones” reaches 0.15, then the average number per ticket will be 30 rather than the initial value of 20. The relationship between the rising allele frequencies at each position and the frequency of the “jackpot combination” (that contains a “one” at all 200 positions) is shown in Figure 2. As can be seen from the figure, it is only toward the end of the process that the winning combination appears, where it then rises dramatically in frequency. This pattern can be explained mathematically by saying that the frequency of the winning combination is an extreme power function of the 200 individual frequencies.

**Figure 2.**
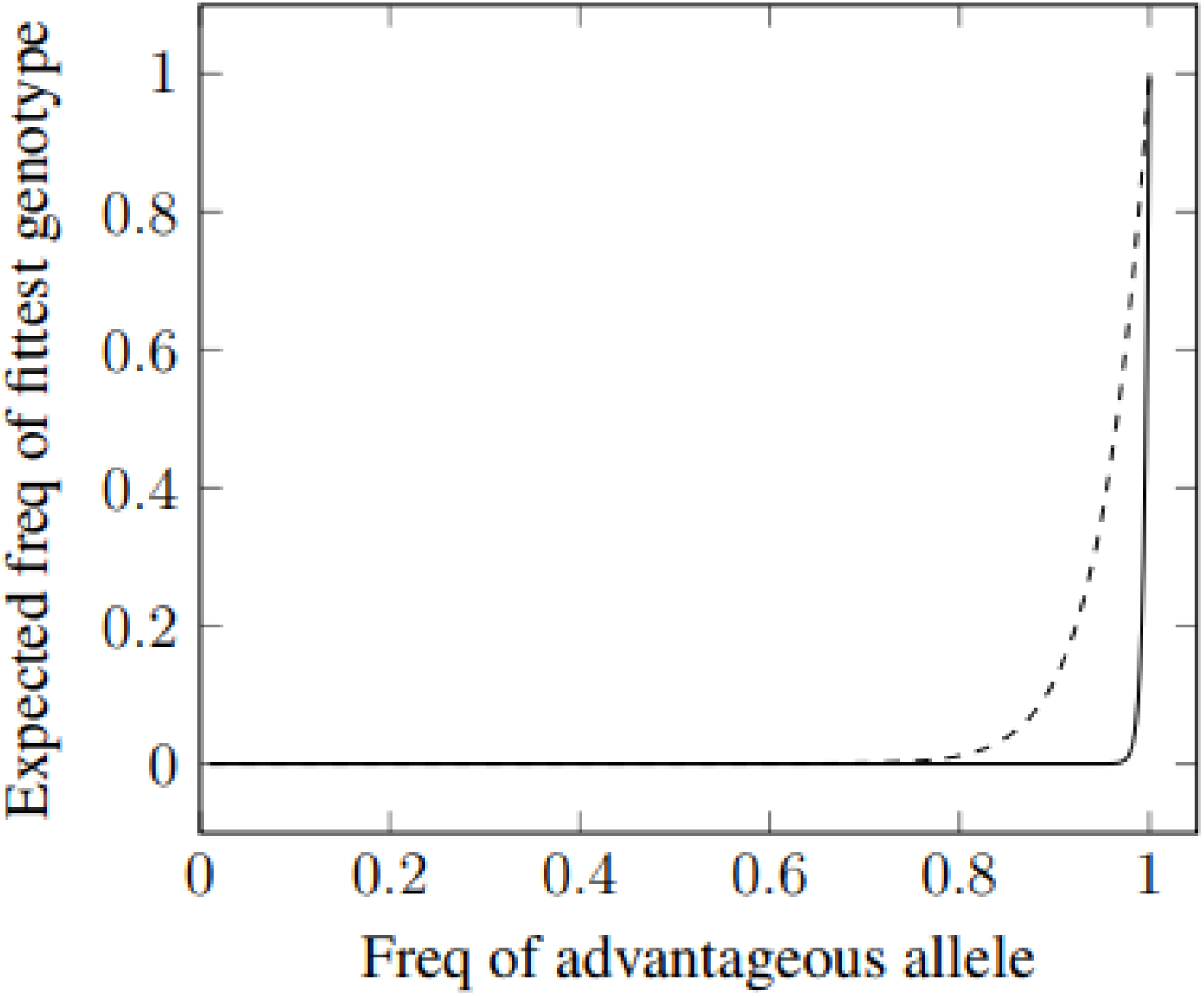
The relationship between the increase in the frequency of favorable alleles at individual genetic loci (X axis) and the expected frequency of the optimal genotypic combination.

As can been seen from the figure, the frequency of the optimal genotype stays close to zero over most of the frequency range at the individual loci. This effect is most pronounced for 200 gene loci (solid line) and less so for 20 loci (dashed line).

In successive rounds of a conventional lottery, new tickets are printed using the same expected frequencies of numbers as in the previous round. But in the genetic lottery these frequencies are modified based on the genetic composition of the selected parents. For example, early in the process the frequency of ones at individual positions may have risen from 0.1 to 0.105 due to selection. In this case random recombination will generate new “tickets” with an average of 21 “ones” per ticket, a slight increase from the original value of 20. As the process continues, both the frequency at individual positions and the total number of ones per ticket will increase concomitantly until, eventually, all tickets will contain the winning combination. In a conventional lottery, such a continual adjustment of the frequencies would be considered a blatant case of cheating. But it is precisely this “trick” that nature uses to arrive at a genotypic combination that appears to be mathematically unattainable at the outset. This process is similar to the multiplicative weights update algorithm, MWUA, that has been used as an optimisation method in game theory and machine learning (Arora et al 2012, Chastain et al 2014). Specifically, random recombination updates the distribution of “tickets” each generation based on the post-selection frequencies of the zeros and ones at each of the 200 positions.

If we restate the problem in more formal genetic terms, we can consider the “tickets” as genotypes and the zeros and ones as allelic variants. At the start of this process, the frequency of the favorable allelic variant at each locus is 0.1. Consequently, the probability of finding one individual with the favorable allele at every locus in the first generation is only 10^−200^. Although the prospect of finding an individual with the optimal genetic combination (i.e., a favorable allele at all 200 loci) is essentially zero at the start of the process, inheritance provides a feedback loop between generations that continuously improves the odds. All genotypes in the population will have a relatively low fitness initially, but natural selection will tend to favor “the best of the bad lot”. These early rounds of selection will result in incremental increases in the frequency of favourable alleles at each of the 200 loci. Recombination will then generate new random combinations of these alleles. And since the individual allele frequencies have increased slightly, the expected frequency of genotypes with higher numbers of favorable alleles per chromosome will also increase automatically. In this way, despite its random nature, recombination facilitates the production of genotypic combinations that would be hopelessly improbable in its absence.

## Discussion

The exponentially increasing number of genotypic combinations as the number of genetically variable loci increases was first pointed out by East (1918). Subsequently, Iles et al (2003) used this relationship to point out that multi-locus genotypes could be very rare even in very large finite populations. Our focus here is on the situation where the number of loci is of the order of 100 or greater. As illustrated in Figure 1, once the number of loci under selection exceeds twenty or thirty, the number of possible genotypic combinations becomes much greater than most biological population sizes. This means that natural selection cannot act on the full range of genotypes. Rather it is limited to a narrow slice of such genotypes in any given generation. It is tempting to think of natural selection as always favoring the fittest possible genotype. But when many loci are under simultaneous natural selection, the theoretically fittest genotype is often absent from the population. Thus, selection cannot act directly on this genotype, although we tend to think of the process in such teleological terms. In fact, the production of the optimal genotypic combination is simply the automatic consequence of recurrent rounds of natural selection and recombination. Most of the selection occurs among genotypes of varying degrees of intermediate fitness. And the effects of recombination are purely random. Yet the interaction between selection and random recombination provides an efficient mechanism for the eventual production of the optimal genotype. The power of recombination is that it allows a finite population to effectively resample genotypes from a virtual infinite population with the current allele frequencies (Ewens 1972, Edhan et al 2017, Hickey and Golding 2018). It is in this way that it produces higher fitness genotypes that were not previously present in the population.

Our argument is based on the assumption that the frequency of any given genotypic combination after a round of recombination can be estimated by multiplying the frequencies at the individual loci. This assumption requires that approximate linkage equilibrium is maintained during the course of the selection. Nagylaki (1993) showed that this is true when the recombination fraction is equal to, or greater than the selection coefficient. For example, two adjacent genes that are one centimorgan apart would remain in approximate linkage equilibrium if the selection coefficient were 0.01. Non-adjacent genes would maintain linkage equilibrium in the presence of higher intensities of selection. This explains the somewhat paradoxical situation that, whereas a given gene can remain in linkage disequilibrium with its nearest neighbors for many generations, it becomes randomized with regard to the majority of other genes rather quickly. Under such conditions, and when the fitness interactions between loci are multiplicative, the multiplicative weights update algorithm can be used (Arora et al 2012). Here, we point out that this process provides an efficient mechanism for the production of higher fitness genotypic combinations that were not previously present in the population.

Previous studies of recombination in finite populations have focused on the adjustment of existing genotype frequencies (Otto and Barton 2001, Iles et al 2003, Otto and Gerstein 2006). Here, our focus is not on the frequencies of existing genotypic combinations, but rather on the production of new genotypes that did not previously exist in the population. As fitter haplotypes increase in frequency, recombination can act to produce still fitter haplotypes (Iles et al 2003). Recombination enables the population to resample the universe of all possible genotypic combinations based on the current allele frequencies (Ewens 1972, Hickey and Golding 2018). Thus, a finite sexual population is not limited to selection among those genotypes that already exist within it. Previously, adaptive evolution based on selection at many genetic loci was seen as potentially very costly in terms of selective deaths (Haldane 1957). However, Haldane’s calculations were based on a model of selection among fixed genotypes. Once we include the possibility of recurrent disassembly and reassembly of genotypes through random recombination, the problem envisaged by Haldane disappears (Hickey and Golding 2019).

This model of selection on the standing genetic variation also points to a possible solution for the problem of the cost of sex that was described by Maynard Smith (1978). This reproductive cost comes from the fact that an asexual female that produced only asexual daughters would be twice as efficient in passing along her genes to future generations compared to a sexual female. This should provide an immediate twofold advantage for a newly arising asexual clone. Such a prediction is indeed true in a non-evolving, monomorphic population. But in a population that is actively evolving based on multi-locus selection, other factors come into play. Most important, the average fitness of the sexual population continues to increase each generation while the asexual clone is genetically frozen. This means that the initial twofold relative advantage is continually discounted as the fitness of the sexual population increases. Simulations have shown that a new asexual clone initially increases in frequency as predicted by Maynard Smith (1978), but this increase is followed by a subsequent decrease in frequency as the fitness of the sexual population “catches up” and overtakes the asexual clone. The asexual clone is eventually eliminated from the population (Hickey and Golding 2018). Thus, the advantage of an asexual clone is short-lived in an actively evolving population.

Previous authors have also compared Mendelian inheritance to a lottery (Williams 1975, Maynard Smith 1978). But the model discussed by those authors was significantly different from the one proposed here. For example, it was assumed that the “winning” genotypic combination would change randomly from generation to generation, thus eliminating the possibility of informational feedback between generations. Neither did they consider the possibility that the number of possible genotypes could be many orders of magnitude greater than the population size. It is worth noting that, according to our model, the production of the optimal genotypic combination is the result of an iterative process that occurs over many generations; it is not the result of a single “lucky toss” as envisaged by Williams (1975). This iterative process can explain why sexual reproduction needs to occur continuously. Peck and Waxman (2000) wondered why intermittent sexual reproduction is not more common in nature and a number of other authors have sought explanations for obligate sex (Kleiman and Hadany 2015, Crouch 2017).

One of the earliest proposals for the advantage of sex was that it generates genetic variants (Weissman 1887). While this is true, sexual reproduction does not usually increase the level of variation relative to that in the previous generation. Rather, it replenishes the genotypic variation that would have otherwise been eroded by selection and random genetic drift. Even more important, it continually shifts the genotypic distribution towards a greater average fitness, as we have shown above. Without sex and recombination, the adaptive evolutionary process would be greatly impeded. Classic population genetics studies (e.g. Haldane 1957) envisaged selection as performing and exhaustive search among all possible genotypic combinations. As the number of combinations explodes in size, however, such an exhaustive search becomes prohibitively difficult. But when we include recurrent rounds of recombination, then natural selection performs an essentially heuristic search that is much more efficient (Hickey and Golding 2018). Through recombination, nature exploits its own form of the multiple weights update algorithm (see Arora et al 2012, Chastain et al 2014). It does this by favoring the best available sub-optimal genotypes and then recombining them. This solution removes the doubts that have been expressed about the application of classic population genetics theory to adaptive evolutionary change (see Messer et al 2016, Reznick 2016). Since recombination allows selection to act independently at each locus, many genes can be selected simultaneously without incurring a prohibitively large reproductive cost.

Our conclusion is that the adaptive function of sexual reproduction lies in its ability to translate increasing allele frequencies at many individual loci into increasing numbers of favorable alleles per chromosome. In the short term, natural selection acts on whole genotypes but recombination then disassembles these selected genotypes and recombines them into new, higher-fitness combinations. In the long term, this two-step iterative process of selection and recombination can produce genotypes containing potentially thousands of advantageous genetic variants. Without recombination, the efficient production of such genotypes would be mathematically unattainable in a finite population. Looking at the problem from another perspective, we could say that recombination is the process that translates short-term genotypic selection into long-term genic evolution.

## Acknowledgements

This work was supported by grant number RGPIN-2020-05733 from NSERC Canada to GBG.

